# Signatures of contextual interference in implicit sensorimotor adaptation

**DOI:** 10.1101/2022.07.03.498608

**Authors:** Jonathan S. Tsay, Carolyn Irving, Richard B. Ivry

## Abstract

Contextual interference refers to the phenomenon whereby a blocked practice schedule results in faster acquisition but poorer retention of new motor skills compared to a random practice schedule. While contextual interference has been observed under a broad range of tasks, it remains unclear if this effect generalizes to the implicit and automatic recalibration of an overlearned motor skill. To address this question, we compared blocked and random practice schedules on a reaching task in which we used a feedback perturbation method that isolates implicit adaptation. The degree of implicit adaptation was quantified as the change in hand angle in the opposite direction of the perturbation, and retention was quantified as the percent of adaptation remaining after visual feedback was extinguished. In two experiments, participants tested under a random practice schedule exhibited slower implicit adaptation, but better retention compared to participants tested under a blocked practice schedule, the signature of contextual interference. These results indicate that contextual interference is not limited to the acquisition of new motor skills but also applies to the implicit adaptation of established motor skills.

## Introduction

Contextual interference is a widely observed phenomena, in which motor skills are acquired faster but poorly retained following a blocked practice schedule compared to a randomized practice schedule (1,2). The ubiquitous nature of contextual interference has come to inform sports instructors and rehabilitation specialists. For example, baseball players who practice hitting curve balls, fast balls, and changeups one skill at a time learned faster but retained less than players who practiced hitting the three types of pitches in a randomized order (3–5). Similarly, patients post-stroke who practiced different compensatory feeding skills in a blocked manner learned faster but retained less than those who practiced the skills following a randomized schedule (6).

Two related hypotheses have been proposed to account for the effect of contextual interference. According to the “elaborative-strategy hypothesis” (1), random practice encourages a learner to compare and evaluate strategies that may be relevant for different motor tasks (e.g., how does preparing for a fastball differ from preparing for a curve ball), and consequently, endows the learner with better contrastive knowledge than that afforded by blocked practice. While the cognitive demands of this exploratory process can produce interference during random practice and, thus, decelerate the rate of learning, randomized practice results in richer and more elaborate long-term motor memories (7,8). Alternatively, the “forgetting-reconstruction hypothesis” (9–11) centers on the idea that random practice results in forgetting between repetitions of the distinct strategies required for different actions (e.g., hitting a fastball or curveball), forcing the learner to continuously reconstruct their explicit strategy with each repetition. While the forgetting process will slow learning, the act of reconstruction will result in stronger long-term memories. Both hypotheses highlight the relevance of strategy and effort during randomized practice that consequently establishes more robust motor memories.

To date, it remains unknown if contextual interference generalizes to the implicit, effortless, and automatic recalibration of an already established motor skill. Consider implicit sensorimotor adaptation of a simple reaching movement, the process by which the sensorimotor system remains exquisitely calibrated in the face of subtle fluctuations in the environment (e.g., a heavier bat) and body (e.g., fatigue after a long baseball game). Given that implicit adaptation places little demands on resource-dependent processes such as decision making and working memory (12–14), one might expect that adaptation might be immune to contextual interference. We are aware of two studies that have tested this hypothesis, one involving force field adaption (15) and the other visuomotor rotation (16), where different perturbations were applied at different target locations. In both studies, subtle signatures of contextual interference effects were observed. However, given the design of each study, it is possible that these effects were related to the recall of explicit strategies (e.g., “aim clockwise from the target on the left”) (17–19), rather than the implicit component of sensorimotor adaptation.

To fill this gap, we used a visuomotor rotation task in which learning is limited to implicit adaptation (Fig 1a) (20,21). On each trial, the participant reached to one of three visual target with the only feedback provided by a visual cursor. The cursor position was time-locked to the radial distance of the hand. However, the angular position of the cursor followed an invariant path, always offset from the target by a fixed angle (“clamped”). Thus, unlike standard visuomotor adaptation tasks, the angular position of the cursor was not contingent on the position of the hand. Despite being fully informed of the manipulation and instructed to always reach directly to the target, participants exhibit a gradual, implicit shift in heading direction in the opposite direction of the cursor. To create conditions for contextual interference, we manipulated the schedule of the three reach locations. For a Blocked schedule, each target was tested in a block of trials, with the three locations tested across successive blocks. For a Random schedule, the three targets were interleaved across trials. We asked if the signatures of contextual interference, namely faster adaptation and worse retention in a Blocked practice schedule, would be observed for implicit sensorimotor adaptation.

**Figure 1.**
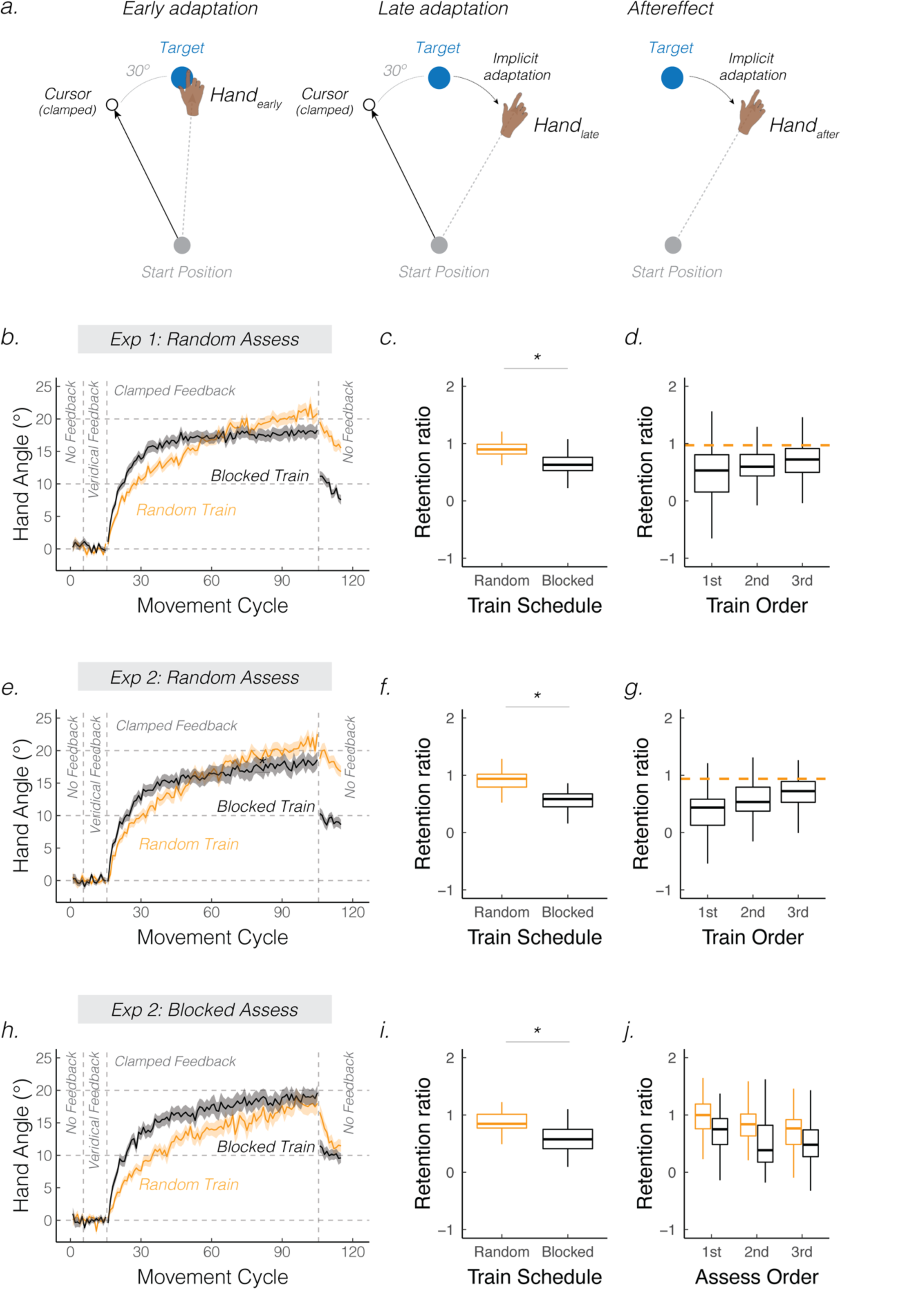
Contextual interference is observed in implicit sensorimotor adaptation. **(a)** Schematic of the clamped feedback task. The cursor feedback (black circle) follows a constant trajectory rotated relative to the target (Exp 1: 30°, 45°, and 60°; Exp 2: 45°), independent of the position of the participant’s hand. Participants were instructed to always move directly to the target (blue circle) and ignore the visual clamped feedback. Left, middle, and right panels display hand and cursor positions during the early, late, and aftereffect phases of adaptation, respectively. **First column (b, e, h):** Mean time courses of hand angle in each experiment. The Blocked Training group is shown in black, and the Random Training group is shown in orange. Shaded error bars denote SEM. **Second column (c, f, i)**: Retention as a function of training schedule. **Third column (d, g, j)**: Retention delineated by the order of targets during the training phase **(d, g)** or by the order of targets during the no-feedback assessment phase **(j)**. Dashed orange line denotes the mean retention over all three targets for the Random Training group. These targets were interleaved, and therefore, do not have a specific order. Box plots show min, median, max, and 1^st^/3^rd^ IQR. Dots denote individuals. * p < 0.05.

## Results

### Experiment 1

During the perturbation phase, participants reached to one of three movement targets, separated by 120°, with a different “clamped” visual error size (30°, 45°, 60°) assigned to each target, counterbalanced across participants. Generalization of implicit adaptation is minimal among targets separated by more than 45° (20,22,23); as such, reaching movements to each target are independently recalibrated. Participants were divided into two groups (N = 120, 60/group): For the Random group, the three targets were interleaved throughout the training phase, and for the Blocked group, the three targets were presented in blocks of 90 trials. For both groups, the three targets were randomly interleaved during the no feedback assessment phase to ensure that retention is assayed in the same way for both groups

The adaptation functions for the Random (orange) and the Blocked (black) groups are shown in Fig 1b. During the baseline phases, participants moved directly to the target. When the clamped feedback was introduced, both groups exhibited a gradual shift in heading direction, approaching an asymptote around 20°, a value convergent with that observed in previous studies that employed the clamped feedback method (20,24). After visual feedback was extinguished, participants exhibited a pronounced aftereffect, a key signature of implicit sensorimotor adaptation. Since this aftereffect was of similar magnitude for all clamp sizes (24), we collapsed over this factor in the following analyses, focusing on the effects of Training Schedule and Phase.

To examine contextual interference, our first analysis compared the two groups at an early timepoint during adaptation (early: first 10 cycles of the training phase) and late timepoint (late: last 10 cycles of the training phase). Participants adapted more in late adaptation compared to early adaptation (main effect of Phase: 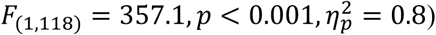, confirming that participants adapted in response to clamped feedback. There was a significant main effect of Training Schedule, with implicit adaptation being on average greater in the Blocked compared to the Random group 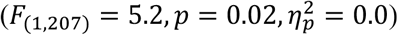. Critically, there was a significant interaction between these factors 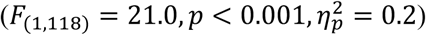: Participants in the Random group adapted less during the early phase than those in the Blocked group (*t*_207_ = 2.3, *p* = 0.04, *D* = 0.7). This difference diminished over the course of adaptation such that late adaptation was slightly larger in the Random group (*t*_207_ = 2.8, *p* = 0.01, *D* = 0.4). Turning to retention, we quantified the magnitude of the aftereffect for each participant by taking the average of their first two cycles of the no-feedback assessment phase and dividing this number by the participant’s late adaptation score (i.e., retention ratio). Using these normalized scores, the Random group showed greater retention than the Blocked group (Fig 1c; Wilcoxon-test: *W* = 2983, *p* < 0.001, *D* = 1.2). Together, these results reveal contextual interference holds for implicit adaptation, namely that a random training schedule results in slower adaptation but greater retention.

However, the Random and Blocked groups have an inherent difference in terms of the delay between training and assessment. For the Random group, the delay between reaches to each target is similar (and small) for the training and assessment phases; that is, the retention test for each target occurs immediately after the end of a training phase that included reaches to all three targets. In contrast, for the Blocked group the delay between training and assessment is substantial for the 1^st^ and 2^nd^ training targets, and minimal for the 3^rd^ training target. Thus, the weaker retention for the Blocked group compared to the Random group (as well as compared to previous studies using the clamped feedback task (21,24)) may reflect the effect of delay rather than the training schedule.

To examine the effect of delay, we honed in on the effect of training order in the Blocked group. As shown in Fig 1d, retention was strongly influenced by delay, being greatest for the 3^rd^ training target (minimal delay between training and retention), and poorest for the 1^st^ training target (largest delay). This result was verified statistically, with the slope of the function relating retention to training order exceeding 0 (robust lmer: *t* = 2.3, *p* = 0.02, *β* = 0.10 *±* 0.04). These results are consistent with previous reports showing that implicit adaptation decays with time between training and assessment (16,25,26).

Given the effect of delay, we performed a stronger test of contextual interference by limiting the analysis to reaches to the third training target location for the Blocked group, comparing retention for this target to the average retention for all Random group. Strikingly, retention remained significantly larger in the Random group (Wilcoxon-test: *W* = 2536, *p* < 0.001). Taken together, we observed marked signatures of contextual interference in implicit adaptation, even when the delay between training and assessment was equalized.

### Experiment 2

In addition to delay, there is a second confound in Experiment 1: The contextual change that occurs in the test phase. For the Random Training group, this change was limited to the removal of the visual feedback. For the Blocked Training group, there was also the change from making repeated reaches towards a single target to reaches towards interleaved targets. The attenuated retention for the Blocked group may reflect, at least in part, an effect of this contextual change.

To address this concern, we adopted a 2 × 2 between-subject design in Experiment 2 (N = 240, 60/group), crossing Training Schedule (Random, Blocked) with Assessment Schedule (Random, Blocked). For the Blocked assessment, there were 10 successive trials for each target. Specifically for the Blocked Training/Blocked Assessment group, the last training target was always assessed first in the aftereffect block to provide a measure of retention of the most recently trained target; for the Random Training/Blocked Assessment group, the target order during the aftereffect block was randomly determined. The clamp size was 45° for all three targets.

We first focus on performance during the perturbation phase. All four groups exhibited robust implicit adaptation (random assessment groups in Fig 1e; blocked assessment groups in Fig 1h). There was a main effect of Phase, with implicit adaptation being greater late compared to the early 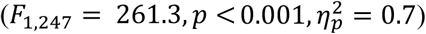. The main effect of Assessment Schedule was not significant 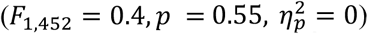, a result that provides a sanity check given that the assessment manipulation does not come into play until the no-feedback assessment phase (and this factor did not interact with the other variables during this phase).

The main effect of Training Schedule was not significant, suggesting that Random and Blocked training groups exhibited a similar degree of implicit adaptation. Importantly, there was a significant interaction between Training Schedule and Phase 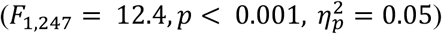: Blocked Training led to faster early adaptation compared to the Random Training (*t*_461_ = 2.8, *p* = 0.02, *D* = 0.7). Similar to Experiment 1, the difference in early adaptation between groups diminished in late adaptation (*t*_461_ = 1.3, *p* = 0.20, *D* = 0.1). However, unlike Experiment 1, we did not observe a significant reversal in learning. (We return to this issue in the following section, “*Pooling together data from all conditions*.*”*)

Turning to retention, we first pooled the data across the first two cycles of the aftereffect phase for each target, ignoring training order and assessment order (Fig 1f & 1i). There was a significant effect of Training Schedule (robust lmer: *t* = 8.6, *p* < 0.001), with Random Training resulting in greater retention than Blocked Training. Critically, the benefit of a random training schedule did not depend on whether the assessment schedule was blocked or random (no effect of Assessment Schedule: *t* = 1.5, *p* = 0.16; no significant interaction between Training x Assessment Schedule: *t* = 2.0, *p* = 0.05).

We then evaluated the effect of delay. Replicating the effect observed in Experiment 1, the degree of retention decreased as the delay between training and assessment increased for the Blocked Training/Random Assessment group (Fig 1f, slope significantly different than zero: *t* = 3.8, *p* < 0.001, *β* = 0.16 *±* 0.04). The Blocked Assessment groups provide a second test of the effect of delay: Retention should decay across the no-feedback phase (i.e., greatest retention for the 1^st^ assessed target, and least retention for the 3^rd^ assessed target). Indeed, retention decreased incrementally with assessment order (Fig 1i, *t* = −3.7, *p* < 0.001, *β* = −0.11 *±* 0.03).

Given the effect of delay, the strongest test of contextual interference requires a comparison of conditions in which the timing of the assessment is roughly equalized following random or blocked training. In the Random Assessment groups, we compared retention for all three targets in the Random Training group to retention for reaches only to the last training target in the Blocked Training group. Correspondingly, in the Blocked Assessment groups, the retention comparison between the two training groups was limited to the first target assessed. Strikingly, retention was greater following random training in both scenarios (Wilcoxon test: Blocked assess, *W* = 2607, *p* < 0.001; Random assess, *W* = 2510, *p* < 0.001). These results highlight a robust contextual interference effect in implicit adaptation, one that holds across different assessment schedules.

### Pooling together data from all conditions

Taking advantage of the large behavioral dataset obtained across these two on-line experiments (N = 360), we pooled the data to examine the overall effect of contextual interference in implicit motor adaptation. As shown in Fig 2, compared to blocked training, random training resulted in attenuated early adaptation (Fig 2a: *t*_340_ = 6.6, *p* < 0.001, *D* = 0.7). By the end of the perturbation phase, adaptation is numerically greater from random training, although there is no statistical difference between the two types of training (Fig 2b: *t*_358_ = 1.9, *p* = 0.06, *D* = 0.2). Most striking is the greater retention associated with random training: Even when limiting the analysis to conditions in which adaptation was immediately assessed, random training resulted in a 17% increase in retention over blocked training (Fig 2c: *W* = 22864, *p* < 0.001).

**Figure 2.**
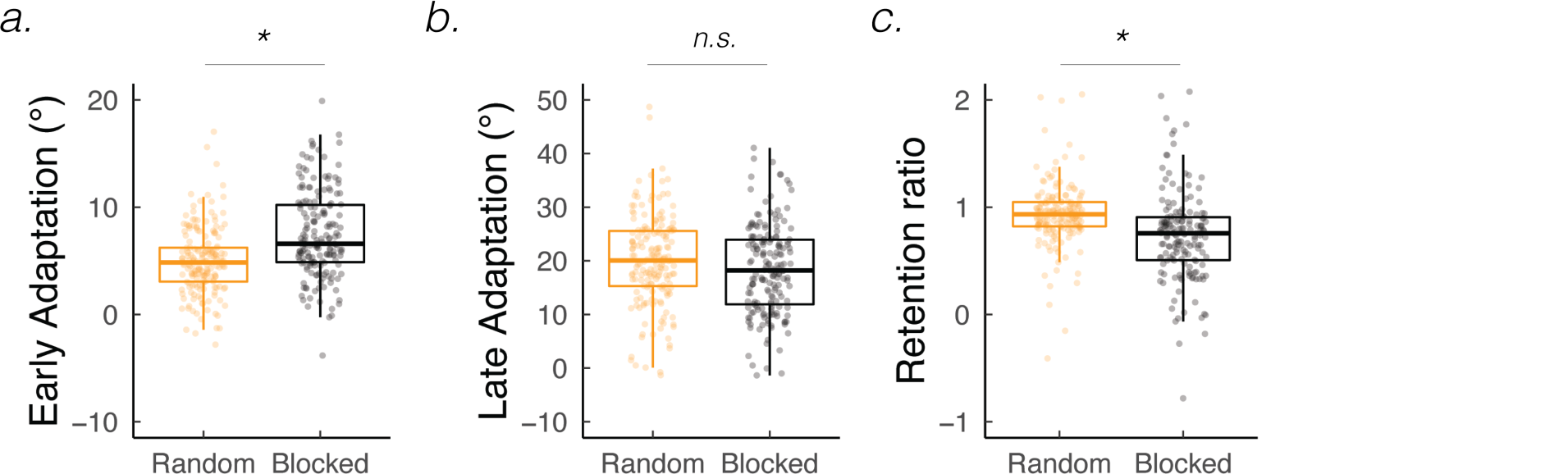
Comparing random and blocked training across all experimental conditions: **(a)** Early adaptation, **(b)** late adaptation, and **(c)** retention. Box plots show min, median, max, and 1^st^/3^rd^ IQR. * p < 0.05. Dots denote individuals (N = 360). Outlier individuals greater than 1^st^/3^rd^ IQR are not shown.

## Discussion

Contextual interference is a widely discussed phenomena in the skill acquisition literature, with random practice schedules resulting in slower acquisition but better retention than blocked practice schedules. Here, we asked whether contextual interference will also manifest during the implicit and automatic adaptation of an established motor skill, reaching. To test this, we employed a visuomotor adaptation task in which performance changes are solely due to the operation of implicit processes (20,21,27–29). In two experiments, we found that participants who performed interleaved reaches to three different target locations consistently adapted at a slower rate but exhibited better retention. These effects persisted even when the schedule of assessment and the timing of assessment were tightly controlled. Taken together, these results broaden the scope of contextual interference to encompass *both* the acquisition of new motor skills and the implicit recalibration of a highly learned skill.

Our findings do not fit easily into the “forgetting-reconstruction” and “elaborative-strategy” accounts of contextual interference. These two hypotheses have focused on how random training enhances top-down control during motor skill acquisition (1,9). As a result, random training imposes greater interference during learning due to the presence of competing strategies, but at the same time, establishes more robust motor memories. However, it is highly unlikely that participants in the clamped feedback task use a re-aiming strategy to offset the visual error (18,30). Not only do the instructions emphasize that they should always aim directly to the target and ignore the visual cursor, but participants also report that their hand position remains near the target throughout the perturbation phase (21). As such, the contextual interference effects elicited in the current studies does not arise from interference occurring during random training between competing (explicit re-aiming) strategies.

A more generic account of contextual interference effects centers on the difference in attentional demands for blocked and random training conditions (31,32). Specifically, while attention to the task is likely to be high near the start of the experiment, it is likely to dissipate as the task becomes familiar. By this hypothesis, the early benefit from blocked practice would come about because the high state of attention allows the system to rapidly come up with a solution. However, over time, blocked practice is likely to lose its attentional hold, leading to reduced retention relative to random practice. While this hypothesis can account for the current results, it is predicated on the assumption that the strength of implicit adaptation is modulated by attentional state. Although the effect of attention on adaptation has been the subject of many studies, this work has generally involved perturbations that engage both explicit and implicit learning processes (19,33,34). Future work using methods that restrict learning to implicit processes would be useful to assess an attentional account of contextual interference.

Another hypothesis of contextual interference in implicit adaptation may be derived from work suggesting that implicit adaptation entails multiple processes that operate at different time scales (35,36). In a two-rate version of this model, one process adapts and decays quickly, operating in the seconds range (“labile” component), whereas a second process adapts and decays slowly, with the effects persistent across days (“stable” component) (25,37–39). We assume that these processes operate in parallel yet are constrained to reach a fixed asymptote due to limits in motor or sensory plasticity (38). As such, they trade-off: If implicit learning is dominated by the labile component, the stable contribution will be reduced. The relative contribution of labile and stable components will differ for blocked and random schedules. Blocked training, entailing repeated reaches to a single target, favors the accumulation of adaptation within the fast, labile process, resulting in fast adaptation but poor retention. In contrast, random training with relatively long temporal delays between reaches to a given target, favors the slow, stable process with the labile component decaying between successive reaches to that target. This would result in slower adaptation yet better retention. Future studies can provide direct tests of this hypothesis, asking how contextual interference in implicit adaptation is impacted by the inter-trial interval between successive reaches. We would predict that the retention cost associated with blocked practice would be eliminated by extending the inter-trial interval.

## Methods

### Ethics Statement

All participants provided written informed consent in accordance with policies approved by the UC Berkeley’s Institutional Review Board. Monetary compensation was provided to the participants for their time.

### Participants

Participants were recruited via two online crowdsourcing platforms, Prolific and Amazon Mechanical Turk. We restricted our recruitment to participants who lived in the United States, had an approval rating greater than 95%, and had completed more than 50 web-based experiments. Participants were excluded if they completed previous web-based reaching experiments sponsored by our lab.

A total of 360 participants were recruited, each of whom completed one experimental session (∼40 minutes). 198 participants identified as male, 146 as female, and 16 as other. Age ranged between 18 – 70 years old (mean ± SD: 32.6 ± 10.7). There were 195 participants who completed the experiment with a computer mouse and 65 participants with a trackpad. We did not enforce any restrictions on device usage since this factor had been shown to not affect reaching behavior and visuomotor adaptation (40).

### Apparatus

Participants completed the experiment by accessing a dynamic webpage created using a customized platform, OnPoint (40). The task progression was controlled by JavaScript running locally in the participant’s web browser. A typical computer monitor has a sampling rate around 60 Hz, with little variation across computers (41). The program automatically detected the parameters of the participant’s monitor and used this information to adjust the size and position of the stimuli. For our sample, the average monitor size was 20-inch with a screen resolution of 1641 pixel width x 940 pixel height. For ease of exposition, the stimuli parameters reported below are based on this average screen resolution.

### Reaching Task

The participant performed reach-like movements by moving the computer cursor with either the trackpad or mouse. On each trial, the participant made a center-out planar movement from the center of the workspace to a visual target. A white annulus (1% of screen height: 0.4 cm in diameter) indicated the center position and a blue circle (1% of screen height: 0.4 cm in diameter) indicated the target location. The radial distance of the target from the start location was 8 cm (40% of screen height). The target could appear at one of three locations on an invisible virtual circle (30°: upper right quadrant; 150°: upper left quadrant; 270°: lower middle).

At the beginning of each trial, participants moved the cursor to the start location. The cursor was represented by a white dot on their screen (0.6% of screen height: 0.2 cm in diameter). When moving to the start location, feedback was only provided when the cursor was within 2 cm of the start location (20% of screen height). After maintaining the cursor in the start position for 500 ms, the target appeared. Participants were instructed that when ready to move, they should produce a fast movement, attempting to “slice” through the target. On feedback trials, the cursor remained visible throughout the duration of the movement and remained fixed for 50 ms at the radial distance of the target when the movement amplitude reached 8 cm. If the movement time exceeded 500 ms or if the reaction time exceeded 2000 ms, the message, “too slow” was displayed in red 20 pt. Times New Roman font at the center of the screen for 750 ms. After each movement, the target (and feedback message when displayed) were blanked and the participant moved back to the start location to initiate the next trial.

### Feedback conditions

There were three types of visual feedback during the experiments: No-feedback, veridical feedback, and clamped visual feedback. During no-feedback trials, the cursor was not visible once the movement was initiated (i.e., when the cursor exceeded 1 cm). During veridical feedback trials, the cursor accurately reflected the participant’s hand position, given the standard horizontal-to-vertical translation associated with manipulating the mouse on a laptop computer. During clamped visual feedback trials, the radial position of the cursor was aligned with the hand, but the angular position was rotated by a constant angular offset from the target (Fig 1a).

### Experiment 1

120 participants were recruited via Amazon Mechanical Turk for Experiment 1. The experiment consisted of 345 trials, divided into four phases (Table S1): A baseline no-feedback phase (15 trials, 5 reaches/target), a baseline veridical feedback phase (30 trials, 10/target), a clamped feedback training phase (270 trials, 90/target), a no-feedback assessment phase (30 trials, 10/target).

Prior to the baseline and assessment phases, an instruction screen was presented informing the participant to reach directly to the target. Prior to the clamped feedback training phase, an instruction screen informed the participant that the cursor would no longer be under their control. The instructions stated that the participant should ignore the visual feedback and reach directly to the target. Six demonstration trials were included to familiarize the participant with the visual clamped feedback. On these trials, the target appeared at 0° (right side of the screen), with clamped feedback provided at a 180° offset from the target. The instructions about the nature of the clamped feedback were repeated before each demonstration trial.

There were two groups of participants (60/group), a Blocked Train/Random Assess group and a Random Train/Random Assess group. Baseline and no-feedback assessment phases were identical for both groups, with the target order pseudorandomized such that each target appeared once every three trials. The key manipulation centered on the structure of the training phase. For the Random Train/Random Assess group, the target location was randomized within cycles of three trials (one/target location). For the Blocked Train/Blocked Assess group, the targets were presented in a blocked fashion: 90 trials for one target, then 90 for the second target, and then 90 for the third target. The order of the targets was counterbalanced across participants in the Blocked Train/Blocked Assess group.

Each target was paired with a single clamp size (30°, 45°, 60°) and the clamp direction (clockwise or counterclockwise) was the same for all three targets. Clamp size, clamp direction, and target location pairings were fully counterbalanced across participants. Note that since contextual interference effects were largely similar across clamp sizes (Fig S1), we collapse across clamp size in the main analyses to focus on the effect of training schedule.

### Experiment 2

240 participants were recruited via Prolific for Experiment 2. Experiment 2 had an identical schedule as in Experiment 1 (345 total trials): A baseline no-feedback phase (15 trials, 5/target), a baseline veridical feedback phase (30 trials, 10/target), a clamped feedback training phase (270 trials, 90/target), a no-feedback assessment phase (30 trials, 10/target). Given the results of Experiment 1, we opted to use a single clamp size (45°) for each target location. The direction of the clamp (clockwise or counterclockwise) was counterbalanced across participants.

We used a 2 × 2 between-participant design in Experiment 2, crossing training schedule (Blocked vs Random) with assessment schedule (Blocked vs Random), yielding four groups (60/group). Random assessment indicates that the three targets were interleaved during the no-feedback assessment phase (as in Experiment 1). Blocked assessment indicates that the three targets were provided in a serial manner. In the Blocked Train/Blocked Assess group, the last training target was always assessed first so that we could evaluate retention of the most recently trained target. In the Random Train/Blocked Assess group, there were no constraints on which target was assessed first since all three targets were learned simultaneously. The order of the three targets during the assessment phase for this group was counterbalanced.

### Data Analysis

The primary dependent variable was hand angle, defined as the angle of the hand relative to the target when the amplitude of the movement reached the target radius (8 cm). Positive hand angle values correspond to the direction opposite the rotated feedback (i.e., we flipped all hand angle values at targets where a counterclockwise rotation was provided). The data were averaged across cycles (three successive reaches), and baseline subtracted to aid visualization. Baseline was defined as mean hand angle over the last 5 movement cycles of the baseline phase with veridical feedback.

Outliers were defined as trials in which the hand angle deviated by more than three standard deviations from a moving 5-trial window, or if the hand angle on a single trial was greater than 90° from the target. These trials were discarded since behavior on these trials likely reflects attentional lapses (average percent of trials removed: Experiment 1: 1.1 ± 0.7%; Experiment 2: 1.4 ± 1.1%).

The degree of implicit adaptation was quantified as the change in hand angle in the opposite direction of the rotation. We calculated hand angle during early adaptation, late adaption, and the aftereffect phase. Early adaptation was defined as the mean hand angle over the first 10 movement cycles during the perturbation phase. Late adaptation was defined as the mean hand angle over the last 10 movement cycles during the perturbation phase. Aftereffect was operationalized as the mean hand angle over the first two movement cycles of the no-feedback assessment phase. Retention was quantified as the percent of adaptation remaining after visual feedback was extinguished, that is the ratio between the aftereffect and late adaptation scores (i.e., retention ratio = aftereffect divided by late adaptation).

The hand angle data were evaluated using a linear mixed effects model (R function: lmer). Post-hoc pairwise statistical tests were performed using t-tests (R function emmeans). P-values were adjusted for multiple comparisons using the Tukey method. Standard effect size measures are provided (*D* for between-participant comparisons; 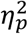 for between-subjects ANOVA) (42). When assumptions of normality were violated, we used the robust linear mixed effects model (R function: rlmer) and the Wilcoxon rank test (R function: wilcox.test), statistical methods shown to be robust to distributional assumptions (43).

## Data availability statement

Data and code will be available upon publication at https://github.com/xiaotsay2015.

